# Tau seeds translocate across the cell membrane to initiate aggregation

**DOI:** 10.1101/2022.05.10.491429

**Authors:** Dana A. Dodd, Michael LaCroix, Clarissa Valdez, Gregory M. Knox, Anthony R. Vega, Ashwani Kumar, Chao Xing, Charles L. White, Marc I. Diamond

## Abstract

Neurodegenerative tauopathies, including Alzheimer’s disease and related disorders, are caused by intracellular aggregation of tau protein in ordered assemblies. Experimental evidence suggests that tau assemblies propagate pathology across brain networks. Tau seeds enter cells through endocytosis but must access the cytoplasm to serve as templates for their own replication. The mechanism by which this occurs is unknown. To study tau uptake, we began with a whole-genome CRISPR knockout screen, which indicated a requirement vacuolar H^+^ ATPase (v-ATPase) components. Treatment with Bafilomycin A1, an inhibitor of the v-ATPase, also reduced tau entry. We next tested direct modifiers of endolysosomal trafficking. Dominant-negative Rab5a expression uniquely decreased tau uptake, as did temporary cold temperature during tau exposure, consistent with a primary role of endocytosis in tau uptake. However, despite reducing tau uptake, these interventions all paradoxically increased intracellular seeding. Consequently, we generated giant plasma membrane vesicles (GPMVs), which cannot undergo endocytosis, and observed that tau fibrils and monomer translocated into the vesicles, in addition to TAT peptide, whereas transferrin and albumin did not. In every case, tau required binding to heparan sulfate proteoglycans (HSPGs) for cell uptake, seeding, or GPMV entry. These findings are most consistent with direct translocation of tau seeds across the lipid bilayer, a novel mechanism of entry into the cytoplasm.

## INTRODUCTION

Neurodegenerative tauopathies such as Alzheimer’s disease (AD) are characterized by neuronal and glial accumulation of tau protein (Lee et al., 2001). Tau is a thermostable, intrinsically disordered protein that binds microtubules (Mandelkow and Mandelkow, 2012). Anatomic patterns of neurodegeneration in tauopathy in many cases follow defined functional neuronal networks (Braak and Braak, 1991; Braak and Braak, 1995; Seeley et al., 2009). A large body of experimental and observational data suggest that, like a prion, tau pathology propagates unique assembly structures, or strains, between adjacent cells and through neural networks (Vaquer-Alicea et al., 2021). Different strains cause distinct clinical and neuropathological phenotypes in experimental mice, and we have thus proposed that they underlie the clinical and neuropathological variation tauopathies (Kaufman et al., 2016).

Our lab originally observed that tau assemblies, or seeds, are readily taken up by cells to trigger intracellular aggregation (Frost et al., 2009). Uptake begins with binding to heparan sulfate proteoglycans (HSPGs), which activates macropinocytosis and tau internalization (Holmes et al., 2013; Stopschinski et al., 2018). This requires specific HSPG sulfation patterns—HSPG mimetics inhibit tau binding, and genetic disruptions of HSPG modification block both uptake and seeding (Holmes et al., 2013; Stopschinski et al., 2018). Once brought into the cell via endocytosis, exogenous tau must somehow exit the vesicle lumen to serve as a template for conversion of endogenous tau monomer in the cytoplasm. The mechanisms by which this occurs are unknown, and could include vesicle rupture (Falcon et al., 2018; Soares et al., 2021; Ugbode et al., 2019), although our recent work suggests this is not required (Kolay et al., 2022). We have developed methods to quantify tau uptake using flow cytometry (Holmes et al., 2013), and to measure induction of intracellular aggregation based on fluorescence resonance energy transfer (FRET) in engineered “biosensor” cell lines (Holmes et al., 2014). This method was recently refined increasing the sensitivity of FRET aggregation assays using stable expression of the tau (P301S) repeat domain (RD) fused to complementary fluorescent proteins, such as Cerulean and Clover (Hitt et al., 2021). FRET flow cytometry measures induction of aggregation in thousands of cells. We have observed that while ~100% of cells take up tau fibrils, in <10% will tau initiate intracellular aggregation (Kolay et al., 2022). Our recent work suggests that tau seeds have two fates after uptake: trafficking to the lysosome for immediate degradation, or entry into the cytosol, where clearance occurs via the proteasome (Kolay et al., 2022). In the vast majority of cases seeding is not associated with vesicle rupture (Kolay et al., 2022). We have now sought to define mechanisms of tau uptake and seeding beginning with a CRISPR-Cas9 screen in HEK293T cells. Our cell and biochemical studies revealed an unexpected explanation for seed entry into the cytoplasm, in which tau fibrils directly translocate across either the plasma or endosomal membrane to promote seeding within the cell.

## RESULTS

### v-ATPase function is required for tau uptake

We screened for tau fibril uptake with the GeCKO lentiviral gRNA library using CRISPR-Cas9, which targets over 19,000 human genes (Sanjana et al., 2014). HEK293T cells were exposed to the library (MOI = 0.3) and cultured for nineteen days with puromycin selection to ensure stable integration of sgRNA and expression of Cas-9. Cells expressing a specific sgRNA were incubated with Alexa 647-labeled, sonicated recombinant 2N4R tau fibrils for four hours. Under these conditions 95.5% of cells took up aggregates. Fluorescence activated cell sorting (FACS) was used to isolate sgRNA containing cells without Alexa 647 staining, gating for the bottom 0.5% cells scoring negative for Alexa 647 (Figure 1).

**Figure 1:**
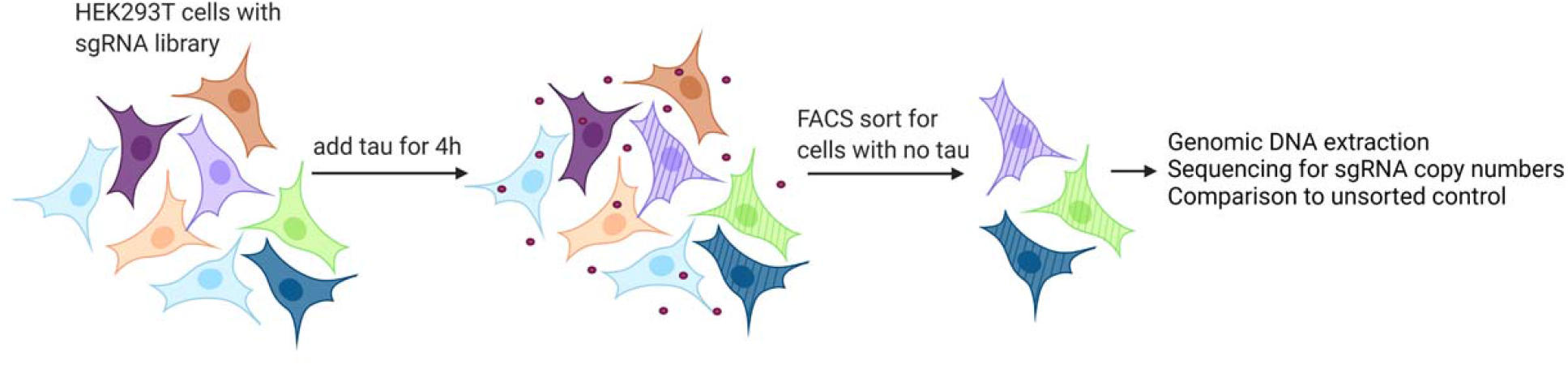
Diagram of the CRISPR screen. HEK293T cells were transduced with the GeCKO sgRNA lentiviral library at an MOI of 0.3. Sonicated, fibrillized Alexa 647-labeled tau was then added to the cells for 4 h. Cells that did not contain Alexa 647 were sorted, the sgRNA was sequenced as a pool, and the copy numbers were compared to the sgRNA sequenced from non-sorted cells. A list of hits is provided as Supplemental Table.

We isolated genomic DNA from Alexa 647-negative cells and from unsorted cells that would represent the full sgRNA library. sgRNA sequences from both samples were amplified and the enrichment of sgRNA in the FACS sorted cells was determined by comparison to the unsorted cells. sgRNA representation was analyzed using MAGeCK (model-based analysis of genome-wide CRISPR/Cas9 knockout). We scored 963 sgRNAs as possible hits (p<0.05) (Supplemental Table) and judged the success of the screen based on the identification of multiple genes associated with HSPGs, including the modifying enzyme NDST1, which we previously determined to be critical for tau aggregate uptake and seeding (Holmes et al., 2013; Stopschinski et al., 2018), and 6 other genes involved in HSPG biosynthesis.

We also identified multiple genes associated with the endolysosomal pathway. The top hit was RNAseK, a vacuolar ATPase (v-ATPase)-associated factor. v-ATPase is a multiprotein complex that hydrolyzes ATP to pump protons across membranes, resulting in the acidification of an endolysosomal compartment. (Perreira et al., 2015). We also identified additional v-ATPase factors, including ATP6AP2, ATP6V0A2, ATP6V2G1, ATP6V1A and ATP6V0D2.

### Genetic inhibition of v-ATPase decreases uptake and increases tau seeding

To confirm RNAseK and ATP6AP2 were required for tau uptake, we synthesized two sgRNAs for each gene, and a sgRNA targeting NDST1, as a positive control that was previously shown to be required for tau binding to the cell surface (Holmes et al., 2013; Stopschinski et al., 2018). We observed lentiviral transduction of HEK293T cells with each of these constructs inhibited tau uptake (Figure 2A). We next tested the effects of v-ATPase gene knockout on tau seeding using a well-characterized biosensor cell line that expresses the tau repeat domain (RD) containing the disease-associated P301S mutation fused to Cerulean or Clover (Hitt et al., 2021). The seed-induced tau aggregation results in FRET that may be quantified by flow cytometry (Holmes et al., 2014). NDST1 knockdown inhibited both tau uptake and seeding. However, knockdown of RNAseK and ATP6AP2 increased tau seeding (Figure 2B).

**Figure 2:**
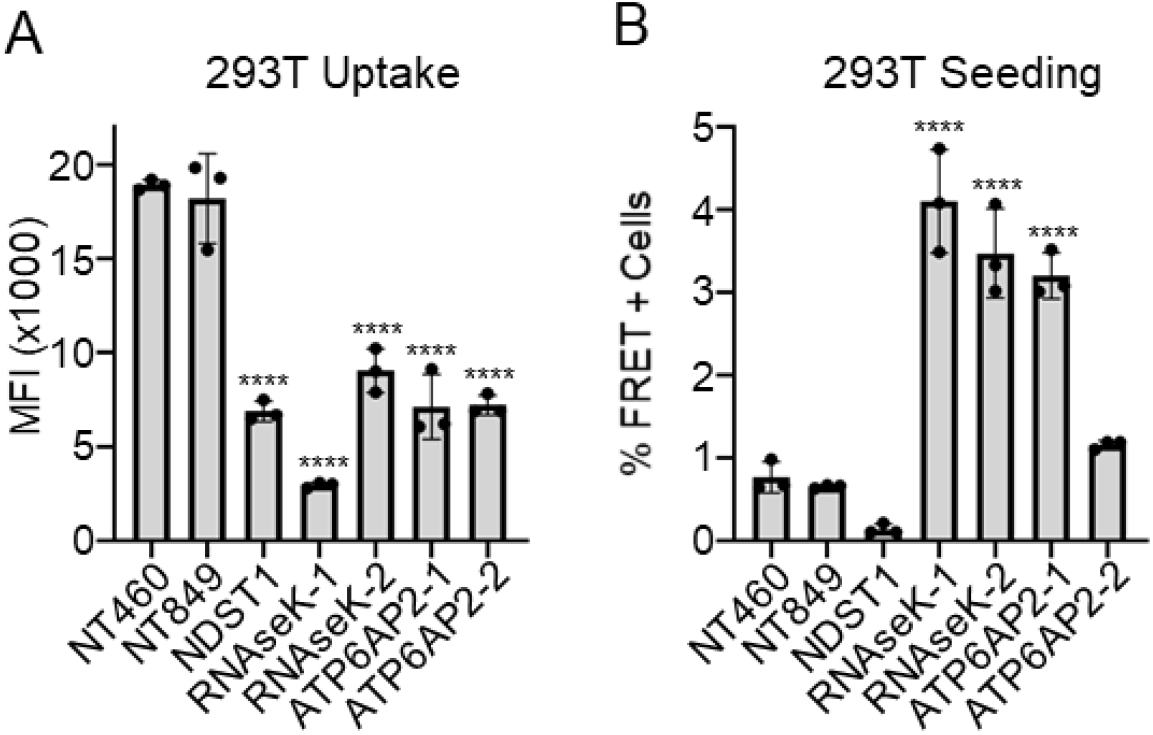
RNAseK and ATP6AP2 knockout reduce tau uptake and increase seeding. A) HEK293T cells were transduced with lentivirus encoding non-targeting sgRNAs (NT460 and NT849) or CRISPR knockout sgRNAs. 25 nM Alexa 647-labeled tau fibrils were incubated with the cells for 4 h. After trypsin treatment, tau uptake was measured by flow cytometry for median fluorescence intensity. This experiment was replicated at least 3 times. B) Tau-RD HEK293T biosensor cells were transduced with lentivirus encoding non-targeting sgRNAs or CRISPR knockout sgRNAs. 50 nM tau fibrils were added to cells and incubated for two days. Cells were harvested and the number aggregate-containing cells was quantified by FRET flow cytometry. This experiment was repeated twice. For both experiments values represent mean ± SD. One-way ANOVA was performed with Dunnett’s multiple comparisons to NT460 with ****=p< 0.0001.

### Bafilomycin decreases tau uptake and increases seeding

Because of our identification of genes involved in endolysosomal acidification as inhibitors of tau uptake, we directly tested this effect by using brief treatment with Bafilomycin A1, which inhibits the v-ATPase by binding a specific subunit (Wang et al., 2021). Bafilomycin is a potent reversible chemical inhibitor of the v-ATPase (Wang et al., 2021). We incubated HEK293T cells with varying concentrations of bafilomycin for thirty minutes prior to exposure to labeled tau fibrils. We harvested the cells at 4 h, and measured tau internalization by flow cytometry. Bafilomycin decreased tau uptake dose-dependently (Figure 3A). To test effects on seeding, we similarly pre-incubated tau biosensor cells for thirty minutes with varying concentrations of bafilomycin and exposed cells for 4 h to tau fibrils and varying concentrations of bafilomycin. We then washed the cells, added fresh media without inhibitor, and incubated cells for 48 h to measure seeding. Seeding increased after the exposure to bafilomycin in a dose-dependent manner (Figure 3B). We next tested this system in iPSC-derived human neurons. We exposed iPSC-derived human neurons to Alexa 647-labeled tau fibrils and 4 h treatment with varying concentrations of bafilomycin, which all inhibited tau uptake (Figure 3C). To measure seeding, we transduced these cells 4 days prior to tau exposure with lentivirus to express tau RD (P301S) fused to Clover or Ruby fluorescent proteins. We incubated cells with tau fibrils and varying concentrations of bafilomycin for 4 h, exchanged the media, and waited 48 h to measure seeding. We observed that all concentrations of bafilomycin increased tau seeding (Figure 3D). We concluded that even brief inhibition of the v-ATPase reduced tau uptake and increased seeding, most consistent with a disruption of vesicle trafficking instead of defects in tau degradation.

**Figure 3:**
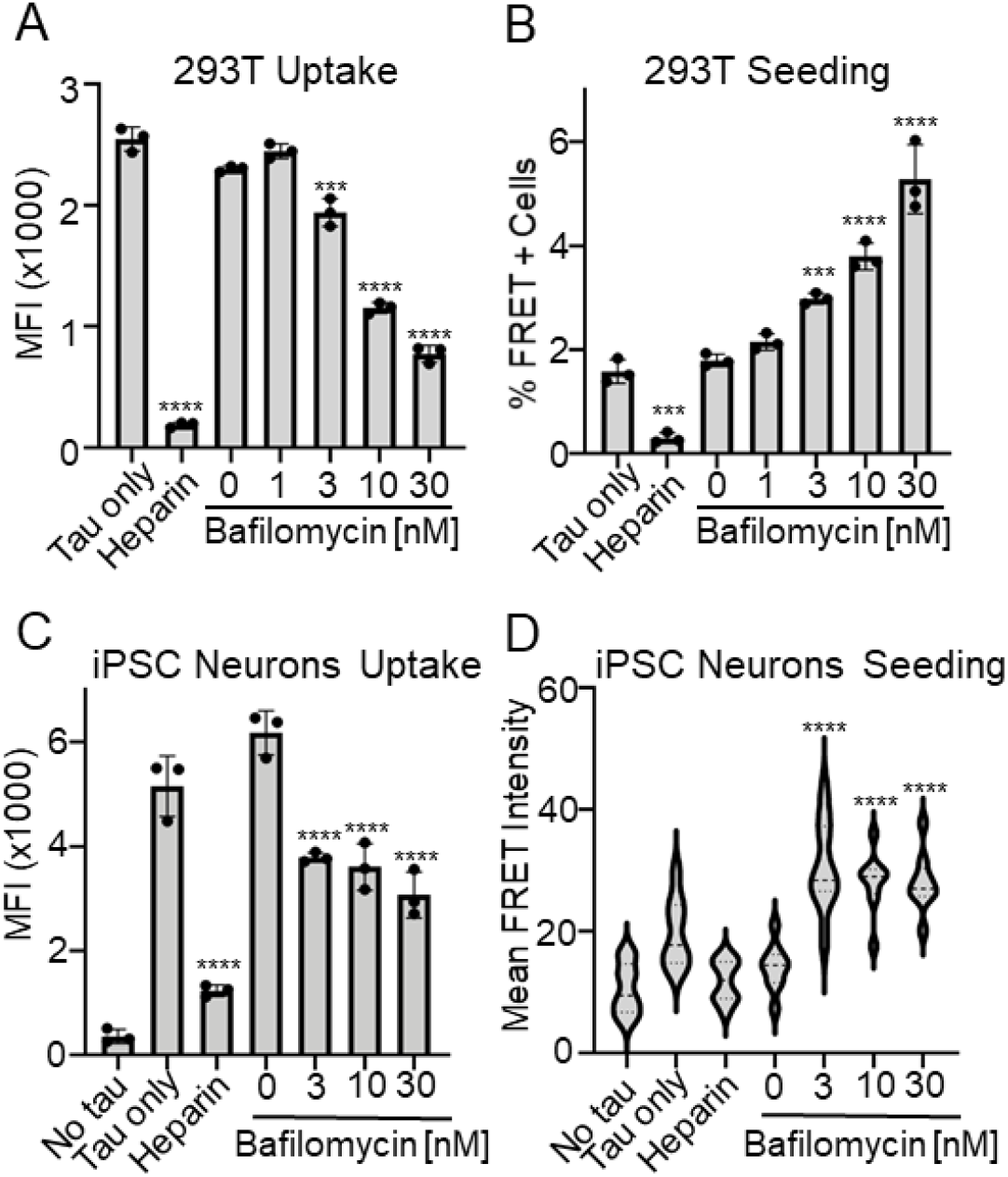
Bafilomycin A1 treatment reduces tau uptake and increases seeding. A) HEK293T cells were pretreated with varying concentrations of bafilomycin for 30 min. 25nM sonicated Alexa 647-labeled tau fibrils were added along with bafilomycin for 4 h. After trypsin treatment, tau uptake was measured by flow cytometry for median fluorescence intensity. This experiment was repeated 3 times. Values represent mean ± SD. One-way ANOVA was performed with Dunnett’s multiple comparisons to 0 nM bafilomycin with ***=p < 0.0005, ****=p<0.0001. B) HEK293T tau-RD biosensor cells were pretreated with bafilomycin for 30 min. Sonicated tau fibrils were added with bafilomycin for 4 h. 200 μg/ml heparin was added as a control. Cells were washed and incubated for 48 h before measuring intracellular aggregation by FRET flow cytometry. This experiment was repeated three times. Values represent mean ± SD. One-way ANOVA was performed with Dunnett’s multiple comparisons to 0nM bafilomycin with ***=p< 0.001, ****=p<0.0001. C) iPSC-derived neurons were treated for 4 h with 25 nM Alexa 647-labeled, sonicated tau fibrils and bafilomycin. Cells were washed and incubated for 48 h before measuring intracellular aggregation by FRET flow cytometry. This experiment was repeated three times. Values represent mean ± SD. One-way ANOVA was performed with Dunnett’s multiple comparisons to 0nM bafilomycin with ***=*p<0.0001. D) iPSC-derived neurons were convertd into biosensor cells by transducing with lentivirus to express tau RD (P301S) Clover and tau RD (P301S) ruby. Four days after transduction 20 nM sonicated tau fibrils and bafilomycin were added. 200 μg/ml heparin was added as a control. Neurons were incubated for 4 h, then carefully washed to remove all bafilomycin, and incubated 48 h to allow seeding to occur. FRET intensity levels were determined by imaging on a confocal microscope with computer analysis. This experiment was repeated three times. One-way ANOVA was performed with Dunnett’s multiple comparisons to 0 nM bafilomycin with ****=p<0.0001.

### Inhibition of vesicle trafficking reduces tau uptake and increases seeding

To test endocytic trafficking more directly, we overexpressed various dominant-negative GTPases linked to different types of endocytosis. Rab5 and Rab7 both control macropinocytosis and clathrin-mediated endocytosis (Buckley and King, 2017; Kimura and Yanagisawa, 2018). Caveolin and dynamin control caveolae-mediated endocytosis, whereas Arf6 and Rac1 are involved in Arf6-mediated endocytosis. Cdc42 controls GRAF1-dependent endocytosis (El-Sayed and Harashima, 2013). Using a lentivirus to express in HEK293T cells the dominant-negative forms of each of these GTPases tagged with RFP, we tested Rab5a(S34N), Rab7a(N125I), Cdc42(T17N), Rac1(T17N), CAV1(Y14F), Eps15(delta 95/295), DMN2(K44A), and Arf6(T27N). Rab5a(S34N) and Rac1(T17N) decreased uptake, whereas Dynamin(2K44A) slightly increased tau uptake (Figure 4A). We next tested the effects on tau seeding and observed that only Rab5a(S34N) increased seeding by approximately 40-fold (Figure 4B).

**Figure 4:**
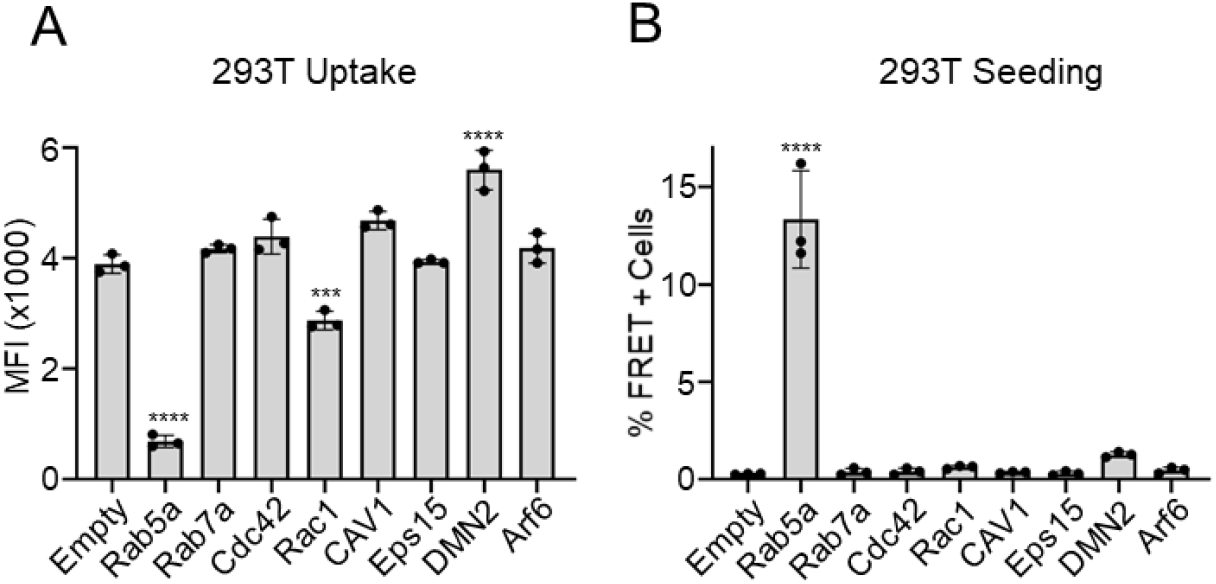
Dominant negative Rab5a reduces tau uptake and increases seeding. A) HEK293T cells were individually transduced with lentiviruses expressing dominant negative proteins involved in different endocytosis pathways. The following dominant negative mutations were used: Rab5a(S34N), Rab7a(N125I), Cdc42(T17N), Rac1(T17N), CAV1(Y14F), Eps15(delta 95/295), DMN2(K44A), and Arf6(T27N). Four days after adding lentivirus cells were incubated with 25 nM sonicated Alexa 647-labeled tau fibrils for 4 h. Tau uptake was measured by flow cytometry for median fluorescence intensity. This experiment was repeated three times. Values represent mean ± SD. One-way ANOVA was performed with Dunnett’s multiple comparisons to empty with ***p = 0.0001, ****p<0.0001. B) HEK293T tau-RD biosensor cells were transduced with lentiviruses that express the same dominant negative proteins as in (A). Four days after adding lentivirus cells were incubated with 50 nM sonicated tau fibrils. After 48 h cells were harvested and intracellular aggregation was measured via FRET flow cytometry. This experiment was repeated 3 times. Values represent mean ± SD. One-way ANOVA was performed with Dunnett’s multiple comparisons to 0 nM bafilomycin with ****=p<0.0001.

### Cold treatment reduces tau uptake increases seeding

The preceding experiments suggested that endolysosomal trafficking defects, while blocking tau uptake, might paradoxically increase seeding. However all interventions had the potential to disrupt protein degradation, even if transiently. To test endocytosis in a different manner, we exploited the fact that it is entirely blocked with temperatures at or below 10°C (Faghihi Shirazi et al., 1982; Tomoda et al., 1989). We temporarily maintained HEK293T biosensor cells at 4° C, 10° C or 37° C for 4 h during exposure to labeled tau fibrils. After treating cells with trypsin to remove any membrane-bound tau, we then either immediately measured tau uptake using flow cytometry, or replated the cells at 37° C for 48 h to allow for seeding. We used heparin as a competitive inhibitor to confirm the requirement for tau binding to HSPGs (Figure 5A).The incubations at 4° C and 10° C blocked almost all tau uptake (Figure 5A).

Despite elimination of tau uptake, incubation at 4° C and 10° C both increased seeding approximately 5-fold. Heparin inhibited tau seeding even at cold temperatures, confirming a requirement for HSPG binding (Figure 5B). To test this mechanism for AD-derived seeds, we immunoprecipitated tau from an AD brain lysate and incubated tau biosensor cells at 4° C or 37° C for 4 hours. Cells were trypsinized, replated, and incubated for 48 h at 37° C to allow seeding to occur. We observed that tau seeding increased with exposure of cells to 4° C when compared to 37° C (Figure 5C). Thus tau seeding induced either by recombinant tau fibrils or fibrils derived from AD patient brain is enhanced when endocytosis is blocked. These findings indicated that tau seeding by recombinant fibrils does not require endocytosis, and suggested that seeds might directly translocate across the plasma membrane after binding HSPG.

**Figure 5:**
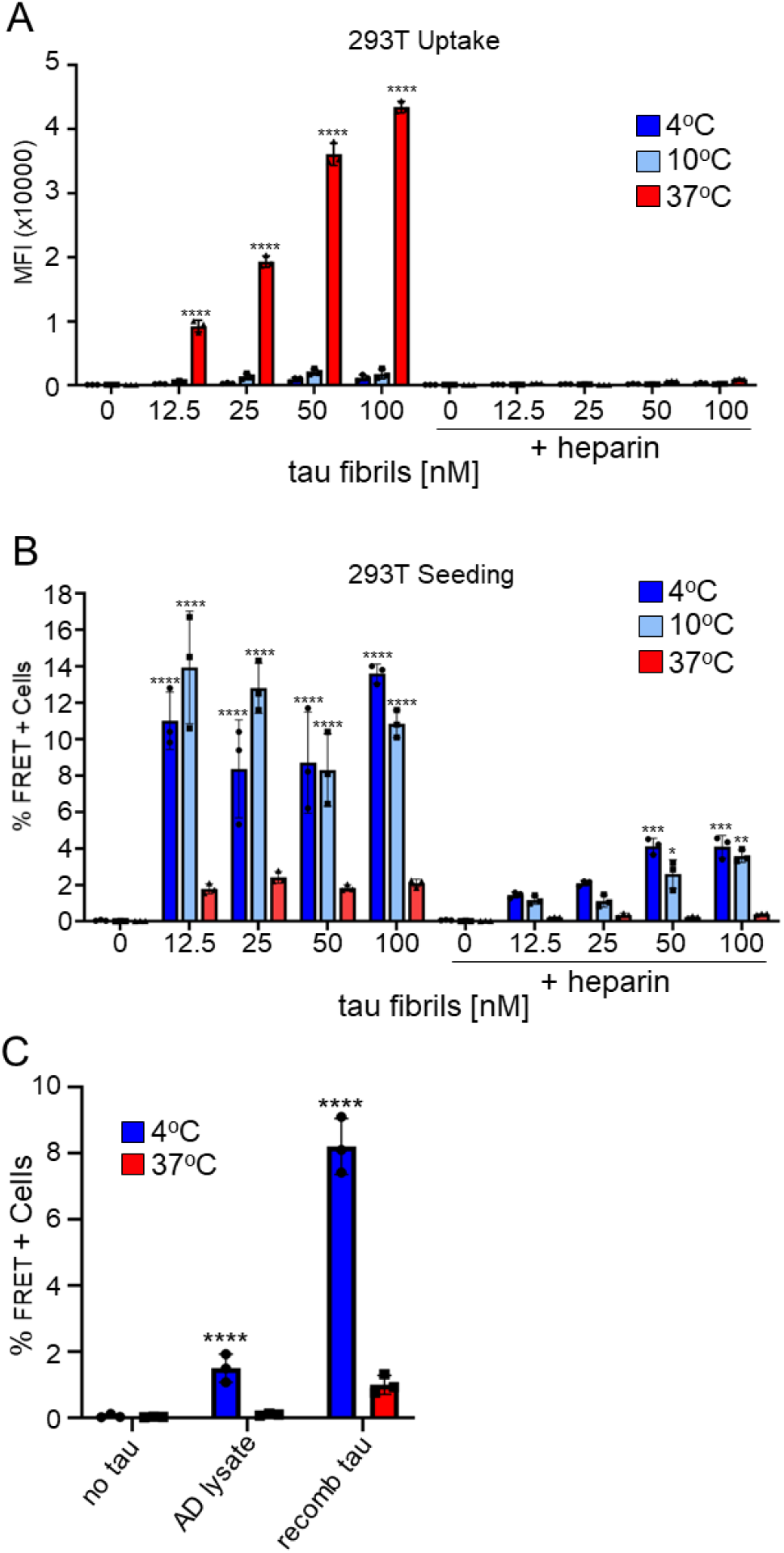
4° C or 10° C incubation reduces tau uptake and increases tau seeding. A) HEK293T cells were incubated with different amounts of labeled, sonicated tau fibrils with or without 200 μg/ml heparin for 4 h while at 4° C, 10° C or 37° C. Tau uptake was measured by flow cytometry for median fluorescence intensity. This experiment was repeated three times. Two-way ANOVA was performed with Sidak’s multiple comparison with ****=p<0.0001. B) Tau RD biosensor cells were incubated with sonicated tau fibrils with or without 200 μg/ml heparin for 4 h at 4° C, 10° C or 37° C. After 48 h cells were harvested and intracellular aggregation determined by FRET flow cytometry. This experiment was repeated 3 times. Two-way ANOVA was performed with Sidak’s multiple comparison with ****=p<0.0001, ***=p<0.0003, **=p <0.002, *=p < 0.04. C) Tau was immunoprecipitated from an Alzheimer’s disease brain lysate and exposed to biosensor cells at 4° C or 37° C. Cells were then trypsinized and replated. After 48 h cells were harvested and intracellular aggregation determined by FRET flow cytometry. This experiment was repeated 4 times. Two-way ANOVA was performed with Sidak’s multiple comparison with ****=p<0.0001.

### Tau translocates into giant plasma membrane vesicles

To test directly whether tau can cross the plasma membrane, and to eliminate the possibility of endocytosis, we utilized giant plasma membrane vesicles (GPMVs) for uptake experiments. GPMVs bud from the plasma membrane of cells upon treatment with 25 mM paraformaldehyde and 2 mM dithiothreitol. GPMVs contain plasma membrane components, including HSPGs, but cannot carry out endocytosis. However, cell penetrating peptides, such as HIV TAT peptide, will directly enter them (Pae et al., 2014). We isolated GPMVs from HEK293T cells and incubated them with 500 μM tau fibrils, tau monomer, albumin, transferrin or TAT peptide. For ease of imaging, we expressed the mCherry fluorescent protein in the HEK293T cells used to form the GPMVs. We labeled sonicated tau fibrils, tau monomer, albumin and transferrin with Alexa 647 while TAT was labeled with FAM. After exposure to labeled proteins at 37° C for 30 min, we washed the vesicles by filtering through spin-columns, before depositing them in 96-well plates for imaging. We used automated imaging (IN CELL, GE) to count thousands of GPMVs in each condition. We determined the percentage of colocalization between mCherry and dye using CellProfiler. Tau fibrils and monomer were readily detected within GPMVs whereas albumin and transferrin did not enter the vesicles. TAT entered the vesicles, as expected (Figure 6A,B) (Pae et al., 2014; Saalik et al., 2011). We also tested for the role of HSPGs in uptake by GPMVs using heparin and heparinase digestion of HSPGs, which has previously been observed to block tau and TAT translocation on cells (Figure 6C,D) (Holmes et al., 2013; Stopschinski et al., 2018; Tyagi et al., 2001). We observed the entry of tau fibrils and tau monomer into vesicles that was inhibited by heparin or heparinase treatment, and therefore concluded that tau binding to the plasma membrane via HSPGs mediates direct translocation, even in the absence of endocytosis.

**Figure 6:**
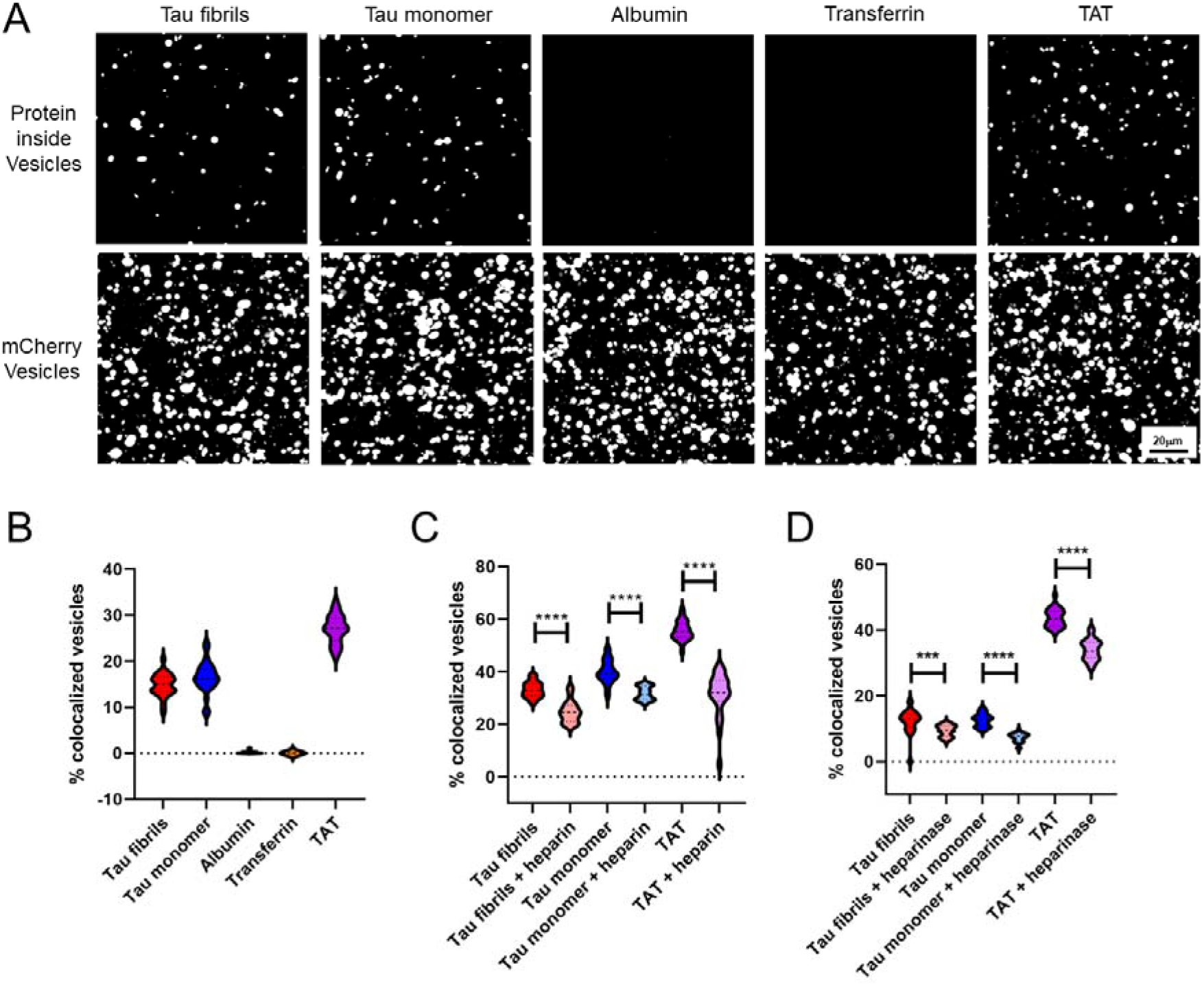
Tau fibrils and tau monomer enter GPMVs via HSPG binding. A) GPMVs were isolated from HEK293T cells expressing mCherry and incubated with 500 nM protein labeled with either Alexa 647 or TAM for 30 min at 37° C. After washing vesicles using a 0.45 μm filter, samples were placed in a glass bottom 96-well plate. Images of vesicles were taken with an InCell analyzer. This experiment was repeated 3 times. B) Twenty-five images from each protein sample were quantified for percentage of colocalized fluorescent signals using CellProfiler. These experiments were repeated three times. C) 500 nM protein was incubated with or without 200 μg/ml heparin for 30 min at 37° C before adding to vesicles for 30 min at 37° C. 25 images from each sample were quantified for the percentage of vesicles with colocalized fluorescent signals. These experiments were repeated 4 times. Comparisons were made by unpaired t test ****=p<0.0001. D) GPMVs were incubated with or without 0.45U/ml heparinase I and III for 30 min at 37° C and washed. Labeled proteins were added to vesicles and incubated for 30 min at 37° C. Vesicles were then washed, imaged, and quantitated. This experiment was repeated 3 times. Comparisons were made by unpaired t test ****=p<0.0001, *** p < 0.0008.

## DISCUSSION

Beginning with a CRISPR screen to find genes involved in uptake of tau fibrils, we identified a role for the v-ATPase protein pump, which was confirmed using Bafilomycin A1, a v-ATPase inhibitor. Our top hit for the CRISPR screen was RNAseK, a v-ATPase associated factor that is required for the internalization of a diverse group of viruses, (Carro and Cherry, 2020; Hackett et al., 2015; Perreira et al., 2015; Zhang et al., 2017), indicating that v-ATPase disruption can be linked to endocytosis inhibition. However, we observed a paradoxical effect of increased seeding in the setting of v-ATPase inhibition. Because vesicle trafficking is affected by acidification defects, we directly tested genes involved in this process, and determined that Rab5a, a protein critical for the development of early endosomes, also reduced tau uptake and increased seeding. To assess the role of endocytosis we next used low temperature to temporarily block endocytosis. This reduced tau uptake to almost undetectable levels, and yet also increased seeding. These results implied that tau could directly translocate across the plasma membrane. To test this hypothesis we measured protein translocation using GPMVs. Tau and TAT peptide each entered GPMVs via an HSPG-dependent mechanism. We concluded that upon binding HSPG on the cell surface, under ordinary conditions tau is rapidly internalized via macropinocytosis, but a small number tau seeds can either directly translocate across the plasma membrane or the vesicle membrane. Under normal conditions, most vesicular tau will traffic efficiently through the endolysomal pathway, with consequently inefficient cytoplasmic seeding (Figure 7). By contrast, we observed that under conditions where vesicular trafficking is impaired, tau seeds have a greater opportunity to translocate into the cytoplasm across the plasma or vesicular membranes.

**Figure 7:**
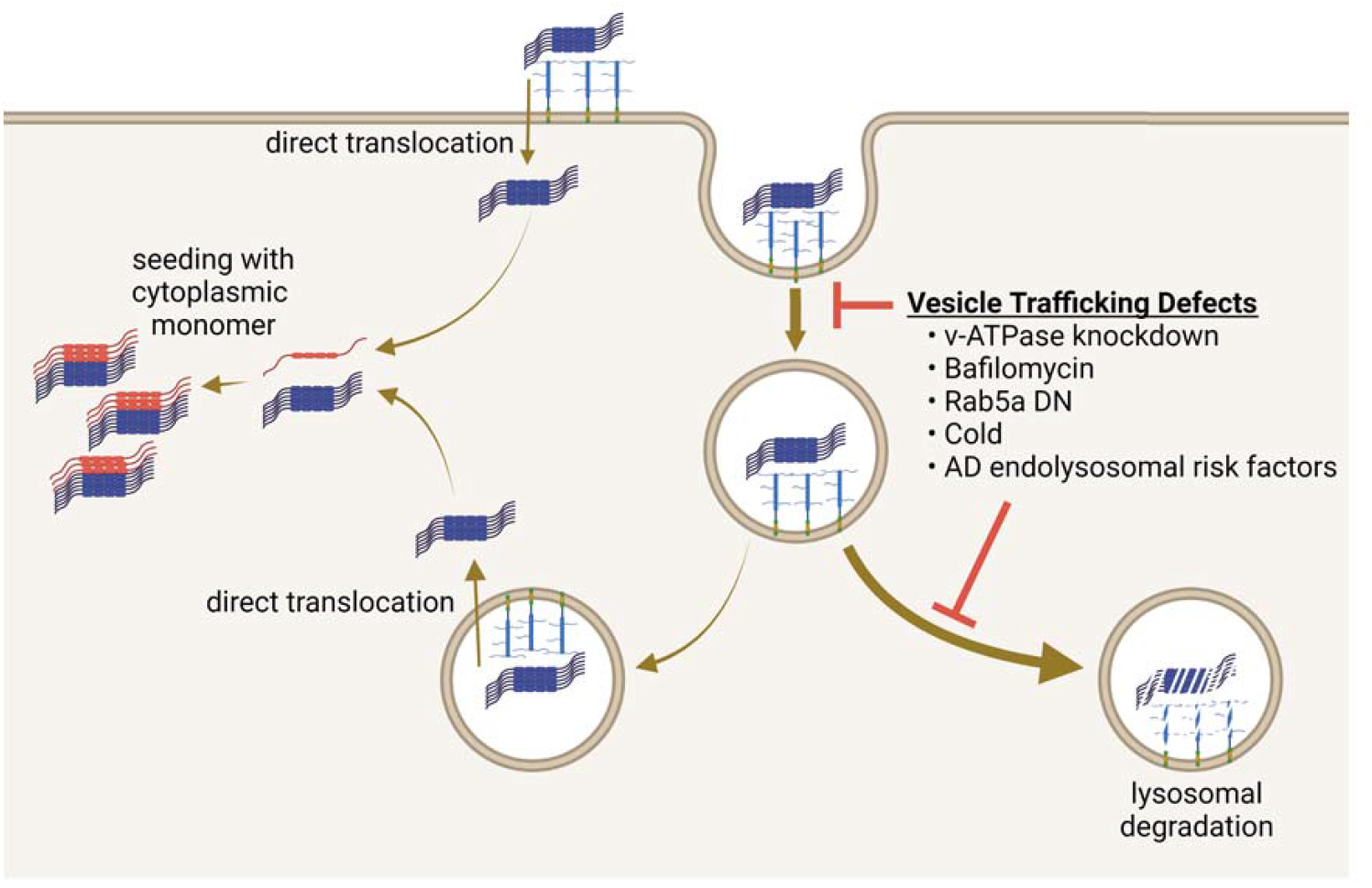
Model for tau entry into cells. Tau fibrils bind to HSPGs on the cell surface. When the cell is functioning properly, most HSPG-bound tau triggers macropinocytosis and trafficking to the endolysosomal system. A small proportion of tau seeds may directly translocate across the plasma endosomal membrane to gain access to the cytoplasm and serve as template to recruit endogenous tau monomer. When the endolysosomal pathway is compromised, increased membrane translocation occurs, with a concomitant increase in seeding within the cytoplasm.

### Tau uptake and seeding are separable processes

Our results indicate that macropinocytosis of tau seeds, which occurs after binding of assemblies of 3 or more tau molecules to HSPGs on the cell surface (Mirbaha et al., 2015), is separable from tau seeding. We learned that disruption of vesicle acidification, even transiently, blocked tau uptake and paradoxically increased seeding. We hypothesized that this occurred because of disrupted vesicle trafficking due to acidification defects. This idea was supported by finding that dominant negative Rab5 and transient hypothermia duplicated the phenomena. In fact, we observed that cold treatment of cells reduced tau uptake to virtually undetectable levels, yet strongly increased tau seeding. These experiments indicated that trapping tau at the plasma membrane, for which HSPG binding was required, actually facilitated seeding. Taken together with our observation of dual fates for tau seeds that enter the cell through endocytosis (Kolay et al., 2022), we have concluded that most tau seeds that enter the cell traffic to a lysosomal degradation pathway, while a small percentage of tau seeds directly access the cytoplasm via translocation across the vesicular or plasma membranes.

### Endosomal processing and tauopathy risk

Many genes associated with AD risk are involved in endocytosis, including *BIN1, PICALM, CD2AP, EPHA1* and *SORL1* (Karch and Goate, 2015; Szabo et al., 2021). In addition, whole-exome sequencing performed on early-onset Alzheimer’s disease patients found multiple genes involved in endolysosomal transport (Kunkle et al., 2017). PICALM, a protein involved in clathrin assembly and autophagosome formation, reduced tau clearance and increased toxicity in a zebrafish model (Moreau et al., 2014), and reduction of BIN1, a protein involved in endocytosis and endocytic recycling, increased the efficiency of seeding in cell models (Calafate et al., 2016). Indeed, others have suggested that disrupted trafficking within the endocytic pathway is linked to AD pathogenesis (Kimura and Yanagisawa, 2018; Neefjes and van der Kant, 2014). Even before amyloid beta deposits appear in the brain, endocytic pathway abnormalities such as Rab5-positive early endosome enlargement, phagocytic vacuole accumulation and defects in lysosomes have been described (Cataldo et al., 2004; Cataldo et al., 2000; Kunkle et al., 2017). Indeed, a recent report indicates that cholesterol depletion, in association with vesicle trafficking defects, promotes cytosolic entry of tau aggregates (Tuck et al., 2022). Our results add a new mechanistic consideration to the role of vesicle trafficking as mediator of tauopathy via effects on cytoplasmic seeding. Specifically, if tau seeds enter the vesicular compartment but do not traffic efficiently towards degradation, they may preferentially access the cytoplasm through direct membrane translocation.

### Tau translocation across the membrane

Others have previously proposed that tau exits the cell via membrane translocation. Several reports have described fibril release from cells by direct membrane translocation from the cytoplasm to the extracellular space, dependent on HSPGs, although the details of how intracellular tau could bind an extracellular glycoprotein are not clear (Katsinelos et al., 2021; Katsinelos et al., 2018; Merezhko et al., 2018). The general idea of protein translocation across membranes is not new, as cell penetrating peptides (CPPs) and some antimicrobial peptides directly translocate across the membrane to move inside the cell (Kauffman et al., 2015; Scocchi et al., 2011). Many CPPs have repeated lysine and/or arginine residues, while the proline-rich region of antimicrobial peptides reportedly mediates translocation. The HIV TAT peptide enters cells by binding to HSPGs and undergoing endocytosis, or directly translocating across the membrane under conditions of hypothermia, and also can directly enter GPMVs (Ben-Dov and Korenstein, 2015; Jiao et al., 2009; Takeuchi and Futaki, 2016). The mechanism(s) of CPP or antimicrobial peptide translocation is unknown. The findings reported here support the surprising idea that even tau macromolecules can directly translocate across membranes, and link vesicle trafficking efficiency to propensity for tau seeding. This raises important biophysical questions about how membranes can transiently facilitate large molecule translocation, without using ATP, and suggests an alternative explanation, besides endolysosomal degradation, for why genes associated with trafficking have been linked to the risk of neurodegenerative disease.

## Materials and Methods

### Cell Culture

HEK293T cells were cultured in DMEM supplemented with 10% FBS and GlutaMax. Cells were grown in a humidified atmosphere of 5% CO_2_ at 37° C.

### Differentiation and culturing of human iPSC-derived cortical neurons

We utilized the integrated, inducible, and isogenic Ngn2 iPSC line (i^3^N). It was previously shown that expression of the transcription factor neurogenin-2 (Ngn2) induces rapid differentiation of iPSCs into glutamatergic neurons (Zhang et al., 2013). This iPSC line harbors a doxycycline-inducible mouse Ngn2 transgene at an adeno-associated virus integration site 1 (AAVS1) safe-harbor locus, allowing for a simplified differentiation protocol (Fernandopulle et al., 2018; Wang et al., 2017). iPSCs were dissociated with accutase (Sigma, A6964) and plated onto basement membrane extract-coated plates (R&D, 3434-001-02). Ngn2 expression was induced with 2 μM doxycycline hyclate (Sigma, D9891) in KSR media alone with 10 μM SB431442 (R&D, 1614), 2 μM XAV939 (Stemgent, 04-0046) and 100 nM LDN-193189 (Stemgent, 04-0074) (doxycycline hyclate is maintained in all medias going forward). On day 2, cells were fed with a 1:1 ratio of KSR media + SB/XAV/LDN and N2-supplmented neural induction media with 2 μg/ml puromycin (Life Technologies, A1113803). On day 3, cells were fed with N2-supplemented neural induction media. On day 4, cells were dissociated with accutase (Sigma, A6964) and plated onto PDL-(Sigma, P1149) and laminin-(Life Technologies, 23 017-015) coated tissue culture plates. Cells were subsequently maintained with neurobasal media (Life Technologies, 21103049) supplemented with NeuroCult SM1 (StemCell Technologies, 05711) and 10 ng/ml brain-derived neurotrophic factor (R&D, 248-BD-005/CF) until collected.

### Protein preparation, fibrillization and labeling of recombinant tau

Full length tau was subcloned into pRK172. Protein was prepared from Rosetta (DE3) pLacI competent cells using a modified protocol described previously (Goedert and Jakes). *E. coli* were grown in Terrific Broth to an optical density of 1.45 at 600 nm. IPTG at 1mM was added and cells were grown for two hours after which bacteria were pelleted and flash frozen. The frozen pellet was resuspended in 1X BRB-80 buffer [400mM PIPES pH 6.8, 5 mM MgSO4, 5 mM EGTA, 0.1% BME] with 1 mM PMSF, 5U/mL DNase and 1 U/mL RNase. A PandaPlus 2000 French press lysed the cells. The supernatant was boiled for 10 min, centrifuged at 15000 x g for 20 min and filtered through a 0.22μm unit. The filtrate was loaded onto a AKTA FPLC with a HiTrap SP HP 5 mL column. There was a gradient elution with 1X BRB-80 buffer and 1 M NaCl in BRB-80 buffer. Column fractions were run on an SDS-PAGE to determine fractions with tau. Tau fractions were pooled and guanidinium chloride was added to a total of 4 M. Dialysis was performed with 4 L of 1X tau buffer [10 mM HEPES pH 7.4, 100 mM NaCl] overnight with one buffer exchange. Tau was then fibrillized with 8uM tau monomer, 8 μM heparin and 10 mM DTT in 1X tau buffer for three days at 37° C without shaking. To add fluorescence 200μl of 8 μM tau fibrils were incubated with 0.025 mg of AlexaFluor 647 succinimidyl ester dye for 1 h at RT. After quenching for 1 h with 100 mM glycine the fibrils were dialyzed into PBS overnight with one exchange of buffer.

### Genome-wide CRISPR screen for tau uptake

#### Library production

The human GeCKO lentiviral sgRNA library v2 (LentiCRISPR) was obtained from Addgene and plasmids were amplified according to the instructions. Lentivirus for Library A and Library B was prepared separately in HEK293T cells in 10 cm dishes transfected using 30 μl Fugene HD with 5 μg Gecko library, 3 μg psPAX2 and 2 μg pMD2.G in 900 μl Opti-MEM. Media was exchanged after 8 h. At 24 and 48 h after media exchange, the media was harvested and pooled. The lentivirus-containing medium was filtered through a 0.45 μm filter and stored at −80° C. The libraries were titered on HEK293T by testing different volumes for puromycin resistance.

#### Transduction of HEK293T with the GeCKO lentivirus library

Biological replicates were used for each library. HEK293T were plated onto multiple 12-well plates and transduced at a MOI of 0.3 with 8 μg/ml polybrene (Millipore). The following day cells were harvested and plated into a T300 flask. 48 h later cells were harvested, pooled and 1 μg/ml puromycin (ThermoFisher) was added to the cells. Cells were expanded and kept in puromycin for nineteen days. The day before cell sorting cells were split to be collected by FACS and used as an unsorted control.

#### Tau uptake and cell sorting

Alexa-647-labeled recombinant tau fibrils were sonicated for thirty seconds at amplitude 6.5 and diluted into media to a final concentration of 25 nM. After adding tau to the flasks of lentiviral-transduced cells they were placed back in the incubator for 4 h. 0.25% trypsin was added for 5 min and cells resuspended. >2.4 × 10^8^ cells were sorted 2 separate times on a FACS Aria II SORP, once quickly with a fast, non-stringent Alexa-647 gate, pooled and resorted with a slower, more stringent Alexa-647 gate. As cells were being sorted, a complete representative of all cells with 500x sgRNAs were harvested as the non-sorted control, also as biological replicates. Each library, A and B, was collected on separate weeks.

#### Genomic DNA extraction and sequencing

All gDNA extraction was performed with a cell lysis buffer and a phenol:chloroform extraction as described (Golden et al., 2017). The sequencing was performed by two rounds of PCR as detailed (Golden et al., 2017).

#### Bioinformatics analysis

Samples were sequenced on Illumina NextSeq 500 with read configuration as 76 bp, single end. The fastq files were subjected to quality check using fastqc (version 0.11.2, http://www.bioinformatics.babraham.ac.uk/projects/fastqc) and fastq_screen (version 0.4.4, http://www.bioinformatics.babraham.ac.uk/projects/fastq_screen), and adapters trimmed using an in-house script. The reference sgRNA sequences for human GeCKO v2.0 (A and B) were downloaded from Addgene (https://www.addgene.org/pooled-library/). The trimmed fastq files were mapped to reference sgRNA library with mismatch option as 0 using MAGeCK (Li et al., 2014). Further, read counts for each sgRNA were generated and median normalization was performed to adjust for library sizes. Positively and negatively selected sgRNA and genes were identified using the default parameters of MAGeCK. Pathway enrichment analysis was performed using ingenuity pathway analysis tool (IPA, http://www.ingenuity.com)

### Tau fibril uptake assay for HEK293T and iPSC neurons

The day before the experiment HEK293T were plated at 20,000 cells/well in a 96-well plate. iPSC neurons were initially plated at 10,000 cells/well in a glass-like 96-well plate. Tau fibrils labeled with Alexa 647 were sonicated for 30 sec at 65 amp. 25 nM of these sonicated tau fibrils were added to the wells. Plates were incubated at 37° C for 4 h. 0.25% trypsin was added for 5 min to digest tau fibrils on the cell surface. Harvested cells were then fixed and prepared in flow buffer. For tau uptake with Bafilomycin A1 (Sigma) treatment, bafilomycin was added 30 min before tau and left on during the 4 h incubation. Experiments were also done without preincubation of bafilomycin before tau addition and bafilomycin was just as effective at decreasing tau seeding (data not shown).

### HEK293T tau fibril seeding assay

HEK293T stable tau RD-Cerulean and tau RD-Clover biosensor cells were plated at 15,000 cells/well in a 96-well plate. The next day, tau fibrils were sonicated for 30 sec at 65 amp. 50 nM of sonicated fibrils were added to the cells and incubated at 37° C for 48 h. Cells were detached with trypsin, fixed with 2% paraformaldehyde and prepared for flow cytometry. For tau seeding with Bafilomycin A1 treatment, bafilomycin was added 1 h before tau addition and left on for a 3 h incubation. Cells were then washed and incubated at 37° C for 48 h. Cells were detached with trypsin, fixed with paraformaldehyde and prepared for flow cytometry. Experiments were also done without pre-incubation of bafilomycin before tau addition and bafilomycin was just as effective at increasing tau seeding (data not shown).

### iPSC neurons tau fibril seeding assay

iPSC neurons were initially plated at 10,000 cells/well on a glass-like 96-well plate. Cells were maintained for a month in media with BDNF and doxycycline. Laminin (Sigma) at 1 mg/ml was added a few days before the experiment. Lentiviruses expressing tau RD-Clover and tau RD-Ruby under a ubiquitin promoter were added to cells at an MOI of 3 and allowed to express for 4 days. Bafilomycin and 20nM fibrillized, sonicated tau was added to cells for four hours. Cells were then washed and placed back in the incubator for 3 days. Cells were fixed and analyzed on an IN Cell Analyzer 6000 collecting 9 data points for each well with lasers tuned for GFP, dsRed and FRET. A computer program was designed to determine the mean FRET intensity.

### Computer program for FRET analysis of iPSC neurons

#### Background correction

To eliminate image noise in subsequent analysis, all images underwent background correction as follows: a Gaussian filter with a sigma value of 100 was applied to the image and this filtered version was subtracted from the original image.

#### Cell segmentation

To segment cells within an image, we first calculated the image gradient of the Donor channel, using the Sobel method, and normalized this output by the gradient magnitude. We then subtracted the resulting normalized image gradient from the original image in order to enhance cell edges in the image. To remove remaining image noise, we applied a Gaussian filter with sigma value of 5. Last, we calculated the Rosin Threshold from the filtered image and used this threshold value to segment cell boundaries.

#### Estimation of FRET signal

First, to correct for bleed-through from other imaging channels (donor and acceptor), we calculated a linear fit (using the polyfit function) for fluorescence intensity between the channel of interest (FRET Channel) and either the donor or acceptor channel. The slopes from these calculations were then used in the following function to calculate the corrected FRET image.

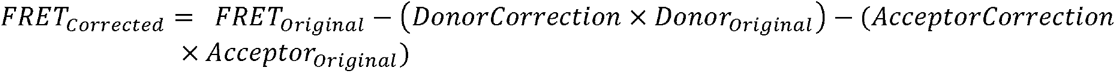

The average FRET intensity for each image was then calculated from the corrected image for all values within the cell segmentation.

### Cold treatment of cells

HEK293T cells were plated as for the uptake or seeding assays on 96-well plates. The next day an Eppendorf Thermomixer was fitted with a plate adaptor and prechilled to 10° C. Samples of tau were prepared in round bottom 96-well plates that were either put on ice (for the 4° C and 10° C samples) or left at room temperature (for the 37° C plate). Plates of cells were then placed on ice or at room temperature while tau samples were added. Plates were then either placed in the 37° C incubator, in the 4° C cold room or on a Thermomixer set at 10° C (not mixing) and for 4 h. Media was removed, warmed 0.25% trypsin added, and cells were placed in the 37° C incubator for 5 min. Warmed media was then added and the cells were gently resuspended. Plates for uptake were then prepared for flow cytometry to quantitate the cells with fluorescence. Cells for the seeding assay were replated and placed in a 37° C incubator for 2 days before being prepared for flow cytometry and FRET analysis.

### Tau immunoprecipitation from Alzheimer’s disease brain lysate

Protein A Dynabeads were washed and 50 μg of Tau B antibody was incubated for one hour at room temperature. Beads were then washed, resuspended in 200 μl of AD brain lysate, and rotated overnight at 4° C. They were then washed and twice with IgG elution buffer (ThermoScientific). Subsequently, Tris buffer pH 8.5 was added. For the seeding assay biosensor cells were plated in a 96-well dish. Either tau-immunoprecipitated AD lysate or recombinant tau fibrils were added to the cells either on ice or at RT. Cells were then moved to 4° C or 37° C respectively for four hours. Cells were then treated with trypsin for 5 minutes, replated, and put in a 37° C incubator for 2 days. Cells were then harvested and counted on a flow cytometer.

### Giant plasma membrane vesicle experiments

GPMVs were prepared according to published protocols with some modifications (Sezgin et al., 2012; Zhao et al., 2016). HEK293T stably expressing mCherry were plated on a PDL coated T300 flask at 2 × 10^7^ cells. The following day cells were rinsed twice with GPMV buffer (1mM HEPES pH 7.4, 2 mM CaCl_2_, 150 mM NaCl) and once with GPMV active buffer (10mM HEPES pH 6.4, 2 mM CaCl2, 150 mM NaCl, 25 mM paraformaldehyde, 1 mM dithiothreitol). Cells were then incubated in active buffer for 6-8h at 37°C. Buffer containing vesicles was collected and left on ice until the following day. The buffer was then concentrated and washed with DMEM FluoroBrite using a 100 kDa Amicon 15 filter tube. When the solution of vesicles was ~2 ml the sample was aliquoted into tubes with 100 μl in each tube. Vesicles with heparinase treatment had 4U/ml of a heparinase I and III blend from Flavobacterium heparinum (Sigma) added for 30 min at 37° C. All samples were then washed with DMEM Fluorobrite using a 0.45 μm filter microfuge. Tau fibrils labeled with Alexa 647 were sonicated at 65amp for 30 sec before using. Heparin treatment of all proteins happened before adding proteins to the vesicles. Proteins had 200 μg/ml heparin added and were incubated at 37° C for 30 min. Albumin and transferrin labeled with Alexa 647 (Invitrogen), TAT(47-57) labeled with FAM (Anaspec), and tau fibrils or tau monomer labeled with Alexa 647 were added to vesicles at a final concentration of 500 μM and put at 37° C for 30 min. Samples were then washed 3 times in a 0.45 μm filter microfuge tube and each sample was put in one well of a 96-well glass plate. Imaging for mCherry, Alexa 647 or FAM was done using an InCell analyzer. 25 spots on each well were imaged. The mCherry levels for the vesicles were variable. To account for all vesicles the brightness contrast was increased until as many vesicles as possible could be detected. There were roughly 100-200 vesicles in each image. The brightness contrast for the Alexa 647 and FAM images were adjusted using mCherry-only vesicles as a guide so that no background was counted. Vesicles were counted using CellProfiler. Primary objects were identified for both the mCherry vesicles and for Alexa 647 or FAM containing vesicles. The object processing module of CellProfiler called “Relate objects”, which can measure whether there is overlap between two fluorescent channels, was used to determine colocalized vesicles of mCherry with either Alexa 647 or FAM. Vesicles that had only Alexa 647 or FAM (a small minority) were not counted. The percentage of colocalized vesicles for all 25 images are represented in the graphs.

## Acknowledgements

Omar Kashmer purified and fibrillized all of the recombinant tau protein used in these studies. Special thanks to Juliane Braun, Josh Mendell and Maikke Ohlson for protocols and advice on the CRISPR screen. Nicholas Rinkenberger in John Schoggins lab kindly provided all of the dominant negative lentiviral constructs. Brian Hitt and Joshua Beaver provided the AD lysate. Sushobna Batra provided the HEK293T cells expressing mCherry. Paniz Karbasi from the BioHPC department contributed a script for image processing. Advice on giant plasma membrane vesicle preparation was provided by Andrea Trementozzi and Jeanne Stachowiak.

